# Int&in: a machine learning-based web server for split site identification in inteins

**DOI:** 10.1101/2023.09.27.559783

**Authors:** Mirko Schmitz, Jara Ballestin Ballestin, Junsheng Liang, Franziska Tomas, Leon Freist, Karsten Voigt, Barbara Di Ventura, Mehmet Ali Öztürk

## Abstract

Inteins are proteins that excise themselves out of host proteins and ligate the flanking polypeptides in an auto-catalytic process called protein splicing. They are gaining momentum in synthetic biology for their ability to post-translationally modify proteins of interest. In nature, inteins are either contiguous or split, in which case the two intein fragments must first form a complex for the splicing to occur. So far, heuristic methods have been employed whenever a new split site in an intein had to be identified. To make the process of split site identification in inteins faster, easier and less costly, we developed Int&in, a web server that uses a gaussian Naïve Bayes machine learning model to predict active and inactive split sites with high accuracy. The model was trained on a data set generated by us and validated using a large diverse data set from the literature, resulting in an accuracy of 0.76. Int&in will facilitate the engineering of novel split inteins for applications in biotechnology and synthetic biology.

## Introduction

Inteins are small intervening proteins translated within host proteins from a single host mRNA. After translation, they perform a so-called splicing reaction to excise themselves out of their flanking external polypeptides (exteins), which are ligated through a new peptide bond (1). Alongside contiguous inteins, split inteins encoded by two separate genes are of particular interest because they allow for a greater variety of applications (2-12) (Fig 1*A*). In the so-called *trans*-splicing reaction, the N-terminal intein fragment (the ‘N-intein’) must first form a complex with the C-terminal intein fragment (the ‘C-intein’). After this step, the splicing reaction takes place resulting in the fusion of the N- and C-exteins (Fig 1*A*). In the past, inteins have been artificially split to obtain either a split intein out of a contiguous one (13, 14), or two intein fragments with low affinity for each other that can be induced to splice when in close physical proximity (14-16). We call a split site allowing the splicing reaction to occur *active*, while one corresponding to two intein fragments unable to splice *inactive*.

**Figure 1.**
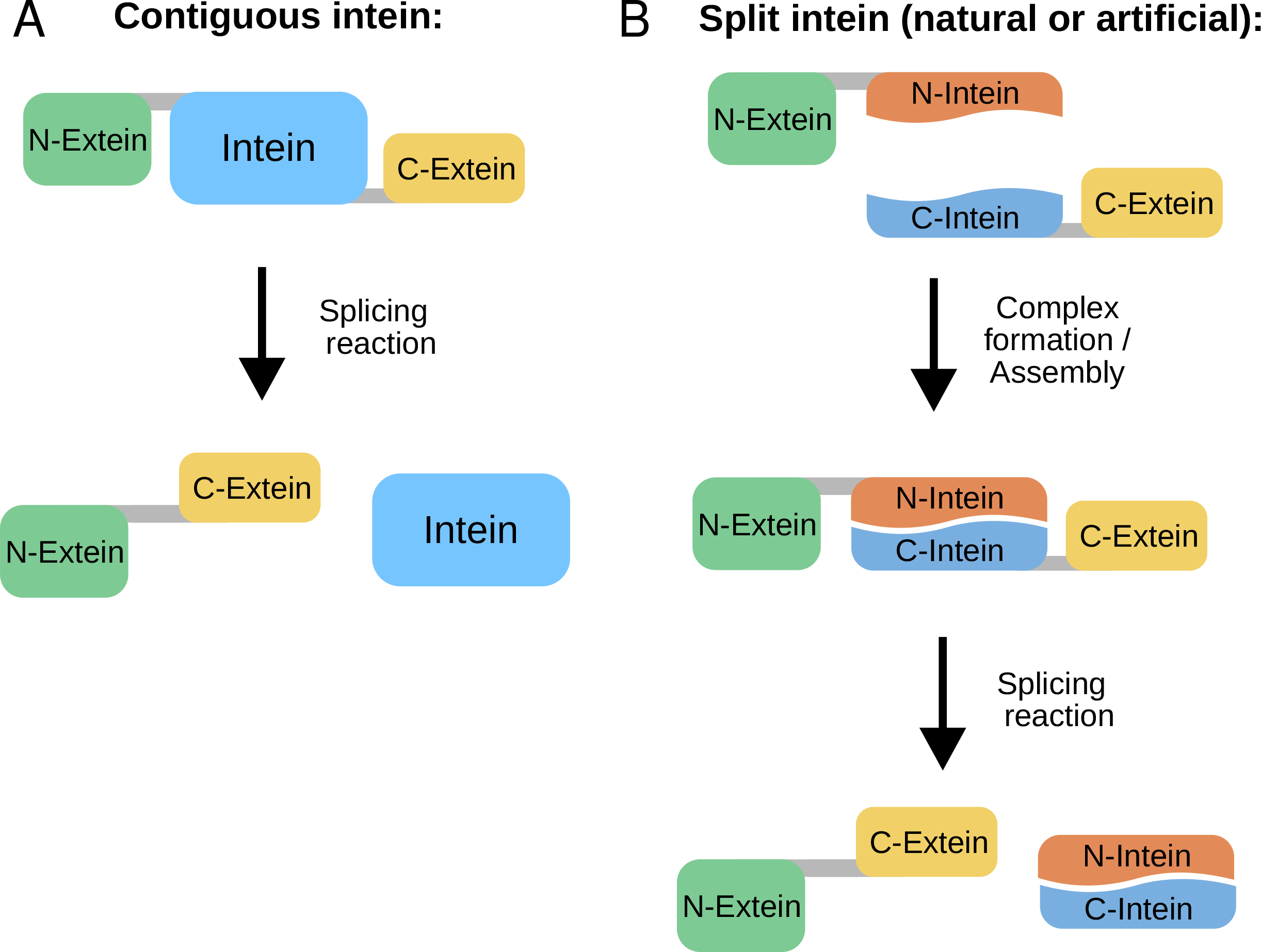
Schematic representation of the protein splicing reaction carried out by contiguous (A) and split (B) inteins.

So far, active split sites in inteins have been found with heuristic approaches (14, 17), or taking very simple protein structure considerations into account (18). The existing SPELL algorithm, that was developed to computationally predict split sites in proteins for the construction of chemogenetic and optogenetic split proteins (19), is inadequate for the specific case of inteins: we found that for ∼80% of inteins collected from the literature (20) no split sites could be predicted (33 inteins without any predicted split sites versus 8 with predicted sites (3D structures were predicted with Alphafold2 (21) as implemented in ColabFold (22); *Appendix*, File S1)). Moreover, for the split sites predicted to be active by SPELL, one site was experimentally shown to be active, while another to be inactive (*Appendix*, Table S1). The ability to identify new active split sites in any intein of choice in an easy and confident manner would represent a breakthrough in the field and encourage a wider range of intein-based applications.

Here we present Int&in, a web server that uses a Naïve Bayes machine learning model to predict active and inactive split sites in inteins with high accuracy. The algorithm predicts the likelihood for each split site to be active as well as the likelihood that the resulting fragments would assemble on their own. We validate the algorithm using data taken from the literature as well as generated by us and show that it has an accuracy of 0.78 for the training set and 0.76 for the validation set.

## Results

### Creation of a data set of active and inactive split sites in inteins

To train the machine learning model, we needed unbiased information not only on active sites –which are easily retrievable from the literature–, but also on inactive ones. Therefore, we decided to generate a data set of split sites using three inteins: gp41-1 (23), *Npu* DnaE (24) and CL (a cysteine-less intein) (25). To cover the whole sequence space of each intein, we randomly selected split sites programmatically with the only rule being that fragments had to be at least four residues long, reasoning that three or less residues would unlikely reassemble with the cognate fragment (26, 27). The split sites were experimentally tested using Western blot as readout with antibodies allowing for the detection of the constructs made of the N-extein and the N-intein (N-construct), and the C-intein and the C-extein (C-construct), as well as the splice product (Fig 2*A*, Appendix, File S2). As exteins we selected proteins known to be soluble in *Escherichia coli*: the maltose binding protein (MBP; N-extein), and thioredoxin and SUMO with a FLAG tag (fused together; C-extein). We tested a total of 126 split sites (36 for gp41-1, 44 for *Npu* DnaE and 46 for CL) (Fig 2*B*). We considered active a site for which a band, albeit faint, could be detected at the size of the splice product. Of the 126 tested split sites, 59 were found to be inactive, and 67 to be active. Efficiency of splicing varied across the split sites (*Appendix* Fig S1, and File S3).

**Figure 2.**
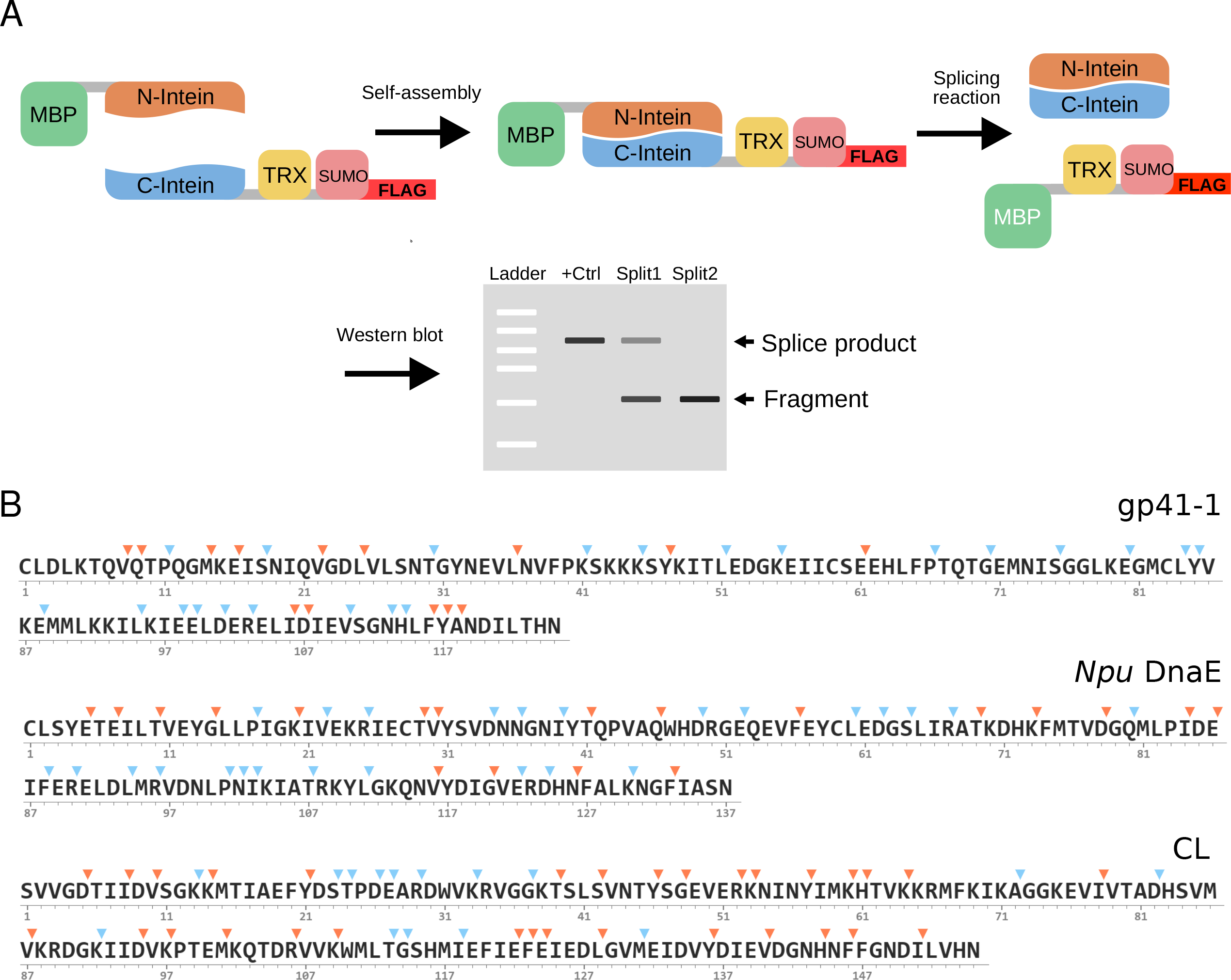
Generation of a split site data set for the development of Int&in. A. Schematic representation of the experimental setup. MBP, maltose binding protein. TRX, thioredoxin. B. Sequences of the inteins used to generate the data set and location of split sites. The color code indicates active (cyan) and inactive (orange) sites as experimentally assessed via Western blot. The numbers indicate residue positions.

### Finding structural and biochemical properties with predictive power

Next, we sought to find structural and biochemical properties within inteins that may be helpful in discriminating between active and inactive split sites. These properties were extracted from the inteins’ structures (for gp41-1 and *Npu* DnaE, crystal structures (PDB id: 6QAZ and PDB id: 4KL5, respectively), while for CL a structure generated through the ColabFold implementation (22) of Alphafold2 (21); *Appendix* File S4). We considered the following properties: i) binding affinity between the two resulting intein fragments, given its strong influence on whether a complex is formed or not (Fig 3*A*) (28, 29); ii) conservation of the residues around the split site, as conserved regions are typically structurally or functionally important and should better not be tampered with (Fig 3*A*); iii) relative surface accessibility of the residues around the split site, considering that residues exposed on the protein surface are likely to contribute less to overall protein stability than those residing in the core, and might thus be a good predictor for active split sites (Fig 3*A*); iv) secondary structure elements, as regions with loops may contribute less to protein structure and could be favorable to locate split sites (Fig 3*A*); v) affinity between the first eight residues of the C-intein and the full N-intein (what we call C-fragment docking) (Fig 3*A*). We considered this property having in mind the ‘capture and collapse’ mechanism introduced by Shah and colleagues, who investigated the naturally split *Npu* DnaE intein through NMR (30). The partially folded N-intein binds to the unfolded C-intein in a process termed *capture*, leading to an intermediate structure formed by the fully folded N-intein electrostatically bound by the still unfolded C-intein. During the *collapse* step, the C-intein folds, generating the functional split intein complex. Assuming generalizability of this mechanism, we decided to consider the C-fragment docking energy as a measure of how easily the intermediate structure of the intein forms.

**Figure 3.**
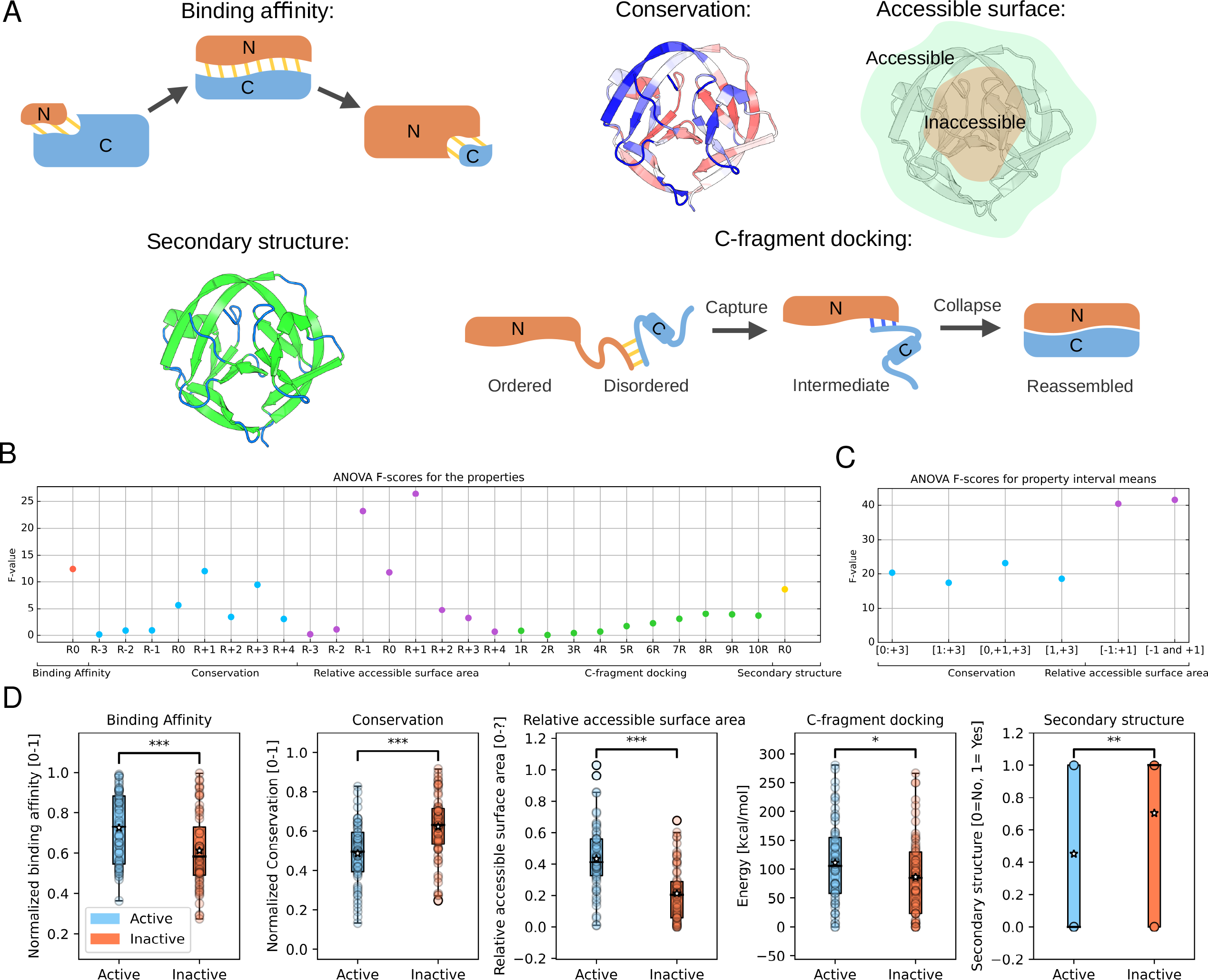
Properties with predictive power for discriminating active and inactive split sites in inteins. (A) Depictions of the different properties investigated. For conservation, conserved residues are shown in red and non-conserved residues in blue. For secondary structure, secondary structure elements are shown in green and unstructured elements in blue. (B) ANOVA F-scores for the different properties. (C) ANOVA F-scores for the mean of several sums of residues’ properties for conservation and relative accessible surface area. (D) Boxplots of active and inactive split sites. A two-sided Mann– Whitney U test was used for all properties but conservation and C-fragment docking for which a two-sided t-test was used. *, p-value < 0.05; **, p-value < 0.01; ***, p-value < 0.001

We calculated the F-value for each property. For the cases including additional residues beyond the split site itself, we calculated the F-value using the mean of the value of that specific property for different combinations of residues (Fig 3*D*). For ‘conservation’, we found that considering residues at positions 0, +1 and +3 gave the highest F-value (Fig 3*D*). Therefore, this combination was finally used for this property. For ‘relative surface area’, residues -1 and +1 were used for the final property. For ‘C-fragment docking’, the highest F-value was obtained when including eight residues.

All the tested properties were able to discriminate, in a statistically significant manner, between active and inactive split sites, being, thus, promising for building the machine learning model.

### Generating the machine learning model

We selected a gaussian Naïve Bayes classifier, which assumes independency of the features it is constructed upon, making it simpler than other algorithms, thus wide-spread in bioinformatic applications (31-33). We tested all possible combinations of properties with a 10-fold cross validation (10 CV) of the Matthews correlation coefficient (MCC) averaged over 10 runs (10x10-fold cross validation) (Fig 4*A*). The combination of conservation, relative accessible surface area, binding affinity, C-fragment docking energy and secondary structure performed best (average 10x10 cross-validated MCC of 0.536, with standard deviation of 0.03), therefore it was used to generate the final model (Fig 4*B*, *Appendix*, Table S2).

**Figure 4.**
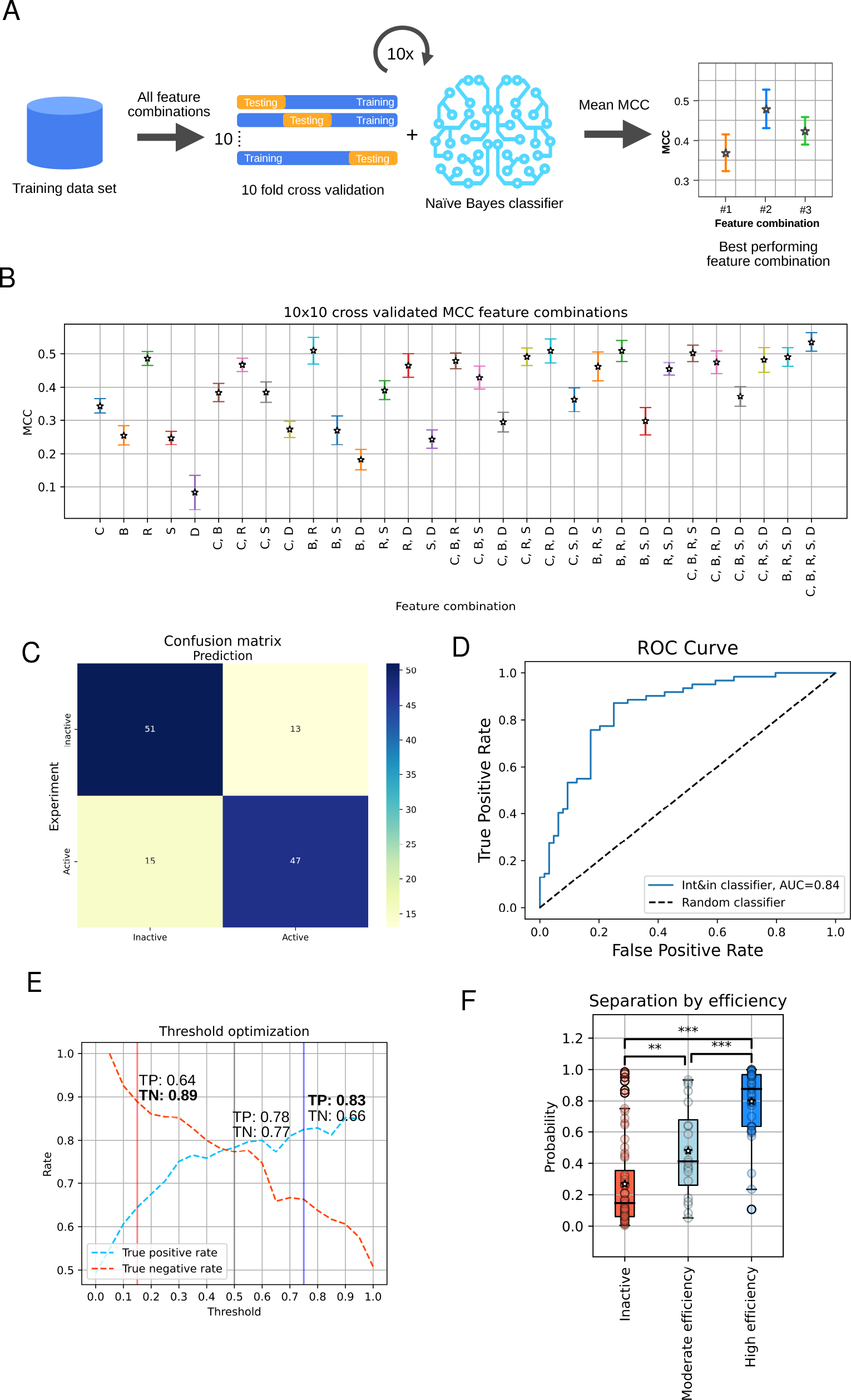
Int&in machine learning model creation with the training data set. (A) Illustration of the workflow for identifying the best feature combination for the gaussian Naïve Bayes classifier. (B) 10x10 CV MCCs plotted with their respective feature combinations. The following letters are used to code for the following features: C = conservation, B = binding affinity, R = relative accessible surface area, D = c-fragment docking, S = secondary structure. (C) Confusion matrix of the Int&in machine learning model with the following features: conservation, relative accessible surface area and secondary structure. (D) ROC curve of the Int&in machine learning model compared to a random classifier. (E) Plot showing the probability threshold adjusted from 0 to 1 with increments of 0.05 plotted against the true positive and false positive rates. The respective rates of true positives are shown for three different thresholds: 0.15 (red), 0.5 (grey) and 0.75 (blue). (F) Plot showing the separation of the inteins according to different splicing efficiencies as predicted by the model. The inactive group shows no activity whatsoever (n=64), the moderate efficiency group contains all split sites with an activity < 50% (n=20) and the high efficiency group contains all split sites with an activity ≥ 50% (n = 42). The two-sided Mann–Whitney U test was used for significance calculation, since the moderately efficient and high efficiency groups are not normally distributed. *, p-value < 0.05; **, p-value < 0.01; ***, p-value < 0.001

With the training data set, the model had an MCC of 0.56, an accuracy of 0.78 (10-fold cross-validated accuracy of 0.77 +-0.13), a precision of 0.78 and a recall of 0.76 (Fig 4*C*). The ROC (receiver operating characteristic) curve, which indicates the performance of a classification model at all classification thresholds in terms of true and false positive rates, had an area under the curve of 0.84 (Fig 4*D*). The model was able to correctly predict 97.5%, 75% and 78.2% of the active split sites for *Npu* DnaE, gp41-1 and CL, respectively (*Appendix*, Fig S2).

The model relies on a threshold to define a split site as active or inactive. Considering that the probability p is a value from 0 to 1, we set the default classification threshold at 0.5, so that any split site with a predicted probability p >= 0.5 is classified as active, while those with p < 0.5 are classified as inactive. With this threshold, the true positive and negative rates are 0.78 and 0.77, respectively. The threshold can be, however, adjusted to specifically increase one of these rates, something that might be required in specific cases (Fig 4*E*). By employing two different classification thresholds, the true positive or true negative rate can be individually increased. As changing the classification threshold also reduces the number of predicted split sites, we set the limit for these thresholds so that the number of predicted split sites does not go below half of the total number of experimental active/inactive sites. Setting the threshold to 0.75, the true positive rate can be increased from 0.78 to 0.83; setting it to 0.15, the true negative rate can be increased from 0.77 to 0.89.

Using these two thresholds, we created four categories of prediction certainty: active split sites (p >= 0.5), inactive split sites (p < 0.5), active with high probability (p >= 0.75), and inactive with high probability (p < 0.15).

Next, we were interested in knowing whether the model could be used to differentiate split sites characterized by lower or higher splicing efficiency (File S2). To this end, the probability of a site to be active, as outputted by the model, was plotted for three groups: inactive split sites, split sites with moderate splicing efficiency (< 50% of the splicing efficiency) and split sites with high splicing efficiency (≥ 50% of the splicing efficiency). We found a significant difference between inactive and low efficiency sites, as well as between moderate and high efficiency sites (Fig 4*F*).

### Model validation

To assess the performance of the model, we extracted from the literature a large data set of 97 split sites in inteins, of which 57 active and 40 inactive, from 41 different inteins (20). We excluded PfuRIR1-1 and PfuRIR1-2, as they were split inside their homing endonuclease/stirrup domain, as well as gp41-1 and *Npu* DnaE, since they are already contained in the training data set. Given the presence of different inteins in this data set, we reasoned it was well suited to evaluate the generalizability of the model, which was obtained with data from three inteins.

We generated the 3D structures of all the inteins with Alphafold2 (21) (implementation in ColabFold (22); *Appendix*, File S1), inputted them into the model, and then calculated performance measures (Fig 5 *A,B*). We found an MCC of 0.51, an accuracy of 0.76 (10-fold cross-validated accuracy of 0.72 +-0.15), a precision of 0.78, a recall of 0.79 and a ROC of 0.83. Applying the same thresholds used with the training data set, the true positive and negative rates could be increased to 0.84 and 0.83, respectively (Fig 5*C*).

**Figure 5.**
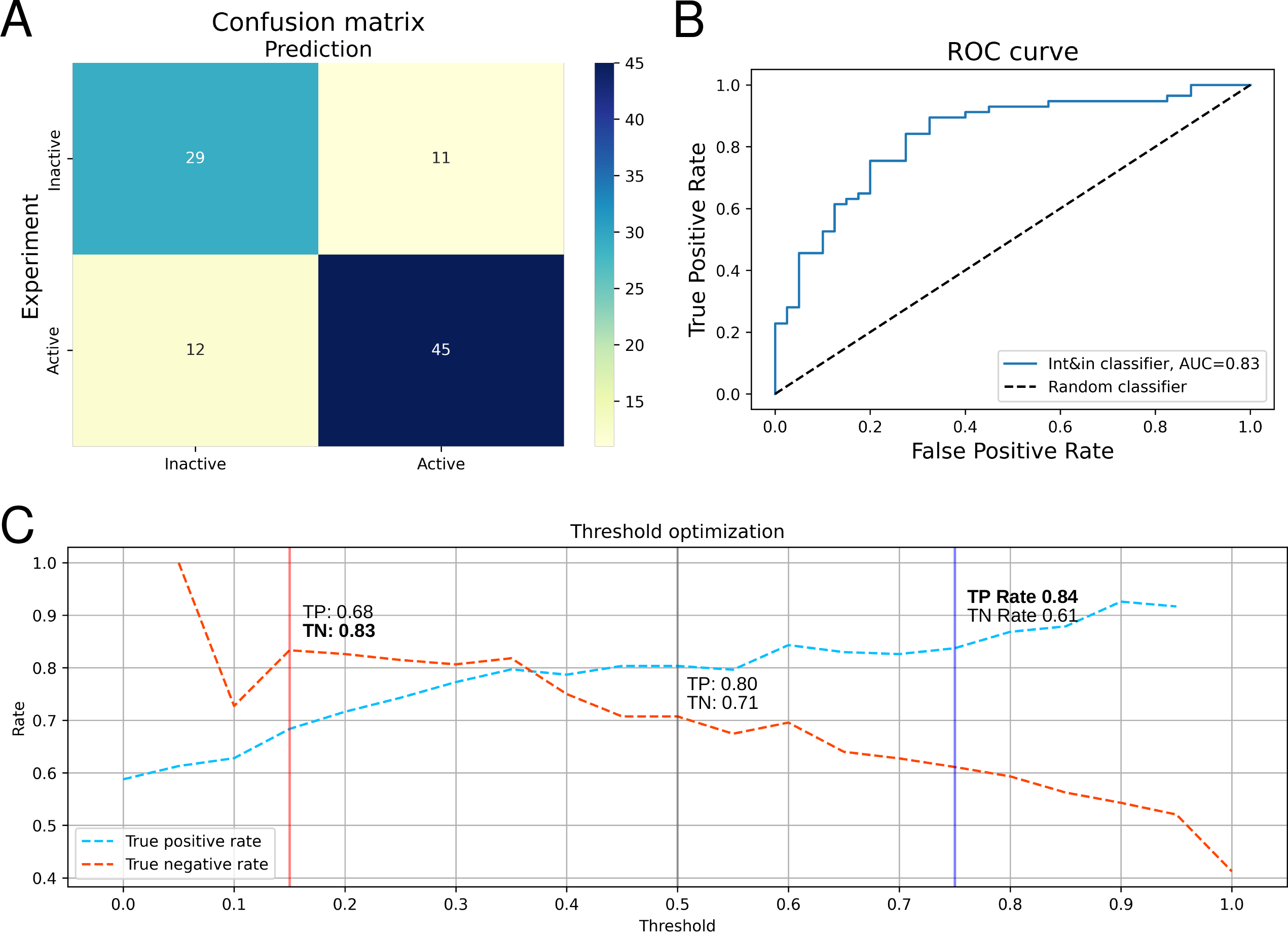
Int&in performs well on a testing data set. (A) Confusion matrix of the Int&in machine learning model with the data set of split sites from literature. (B) ROC curve of the Int&in machine learning model compared to a random classifier. (C) Plot showing the probability threshold adjusted from 0 to 1 with increments of 0.05 plotted against the true positive and false positive rates. The respective rates of true positives are shown for three different thresholds: 0.15 (red), 0.5 (grey) and 0.75 (blue).

### The Int&in web server

To allow researchers of varying level of computational knowledge to make use of this model, we opted for making it available through a web server. The Int&in web server evaluates each amino acid of a given intein sequence for its potential to be an active split site, and provides an easy-to-use web interface to quickly evaluate the results. Users can anonymously upload their experimentally determined or modelled protein structures and receive a personalized link to visualize the results. Additionally, we provide the option to submit batch runs with multiple .pdb files. Finally, the user can register for an account, which ensures that past jobs are easily retrieved and can be analyzed at a later point (past jobs may be deleted after 30 days). After all calculations are executed on the backend, the user is notified via email (if an address was provided at job submission; otherwise, the job can be accessed through the personalized link that appears after job submission), and can access an interface consisting of a structure window, a list of split sites and several graphs showing the values of the individual properties as well as the model prediction. All files generated by the Int&in web server as well as the raw data from third party programs (conservation calculations and secondary structure calculations) can be downloaded.

## Discussion

Here we described Int&in, a web server that relies on a machine learning algorithm to predict active and inactive split sites in inteins. Before embarking in the development of this web server and the model it relies on, we checked the literature and found SPELL, a web server for the prediction of split sites in proteins specialized in the task of protein functional reconstitution by means of ligand-or light-regulated heterodimerizing systems (19). Indeed, locating functional split sites in proteins is a highly desirable task, because splitting proteins into two dysfunctional halves, which can be brought back into close physical proximity to regain activity, is a useful technique in cell and synthetic biology to control and/or understand biological processes (34-36). The algorithm behind SPELL makes use of an energy function and several structural and sequence-based parameters to determine functional split sites. Because it was generated from a limited data set of 27 functional split sites from 16 different proteins, it is difficult to assess whether the rules employed by SPELL truly reflect the full range of sites at which a protein can be split. This was not the main goal of the SPELL algorithm, which aims to maximize the number of true positive sites while keeping the number of false positive sites at a minimum. This means that several functional split sites that are not deemed optimal by the algorithm are lost, which can be problematic when wishing to split a protein in a specific region. For inteins in particular, the possibility to find split sites in specific regions can be crucial to, for instance, generate very short intein fragments, which can be more easily chemically synthesized and used in protein semisynthesis (27, 37, 38) or be more amenable to control via caging within the light-sensitive LOV domain (39). Interestingly, when applied to 41 individual inteins, SPELL predicts no split sites for around 80% of the inteins tested (*Appendix*, Table S1). Additionally, when we applied the same energy profile used by SPELL, which is conceptually equivalent to the binding energy we use, we found no significant difference between inactive and active split sites (*Appendix*, Fig S3). This indicates that SPELL is not particularly suitable to predict active split sites in inteins.

Our model uses the binding affinity, conservation, relative accessible surface area, C-fragment docking energy and secondary structures of and around the split site to calculate a probability score. Being trained on a data set with randomly distributed split sites from three different inteins amounting to 126 sites in total, the model managed to achieve an accuracy of 0.76 with an untrained data set consisting of 41 different inteins with split sites gathered from literature. By dividing the probability scores into four categories (active, inactive, high probability to be active and high probability to be inactive), we managed to further increase, for the testing data set, the rate of true positive or true negative predictions to 0.84 and 0.83, respectively.

As the model was not trained with any knowledge of the efficiency of splicing of each site, but nonetheless managed to output probabilities that show a significant difference between moderately and highly efficient split sites, we speculate that the properties used in the model are also predictive of split site efficiency, and thus Int&in could be used also to predict efficiencies. By testing the model on a data set coming from the literature, characterized by a variety of inteins with different exteins expressed in different model organisms, we have showed generalizability as well as wide applicability of our model. Nonetheless, in cases where the local exteins are changed or truncated the model’s output may show discrepancies, considering the model was trained on results generated with the optimal local exteins for each intein.

Since the Int&in web server is user-friendly, and requires no prior knowledge on bioinformatics or computational biology, we believe it will boost research on these fascinating proteins as well as applications based on them.

## Materials and Methods

### Non-covalent bond determination in Int&in

A list of residues that are in proximity to each other (10 Å at most) is created and the existence of non-covalent bonds (hydrogen bond, salt bridge, van der Waals, pi-pi and pi-cation) is determined, as outlined below.

#### Hydrogen bonds

A hydrogen bond between two atoms is considered to be present if the angle between the hydrogen donor (D) and the hydrogen acceptor (A) is ≤ 63° and the distance between them is <= 3.5 Å (40). N, O, S and atoms may be acceptors and/or donors with the exception of NH_3,_ which cannot be an electron acceptor. Moreover, C atoms may be donors, to account for backbone hydrogen bonds (41). Donor atoms additionally need at least one hydrogen atom covalently bound to them.

#### Salt bridges

A salt bridge between two residues is considered to be present if they have opposite charge and the distance between any of the nitrogen atoms in the side chain of the positively charged residue and the oxygen atoms (or the sulfur of cysteine) in the side chain of the negatively charged residue is ≤ 4 Å (42).

#### Aromatic interactions

*π*-*π* interactions are attributed to aromatic amino acids if the centroids of their aromatic rings are at most 7.5 Å from each other and none or only one of the aromatic rings is positively charged (43). Cation-*π* interactions are considered to be present if the distance between the centroid of the aromatic ring of a neutral aromatic amino acid and any nitrogen atom of the side chain of a positively charged lysine or argine is <= 6 Å (44).

#### Van der Waals

A Van der Waals interaction is considered to be present between any carbon-carbon, carbon-sulfur, carbon-oxygen (glutamine OE1 and asparagine OD1) and carbon-nitrogen (glutamine NE2 and asparagine OE1) if their distance is ≤ 0.5 Å (40). The van der Waals radii (NACCESS radii) of the two potentially interacting atoms is subtracted from the distance between the atoms to get the distance between the atoms’ surfaces (45).

### C-fragment docking energy determination in Int&in

The docking energy between the first eight residues of the C-intein and all residues from the N-intein is calculated by attributing different energies to each non-covalent interaction depending on its type (and distance, in the case of hydrogen bonds):

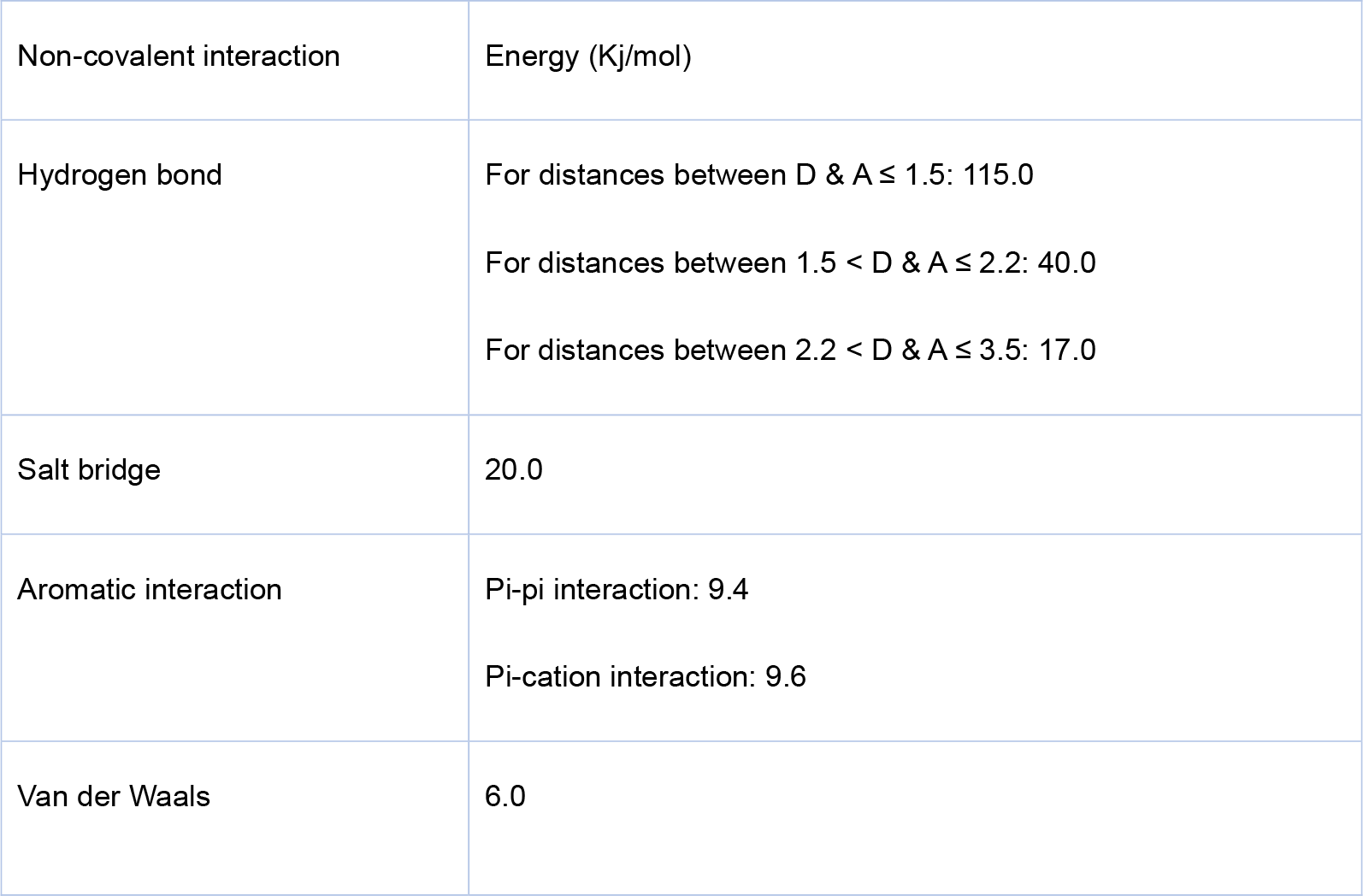

The energies have been used according to the RING 3.0 web server (46).

### Relative accessible surface area determination in Int&in

The accessible surface area of each residue is calculated with the ‘rolling ball’ algorithm by Shrake & Rupley (47). The Van Der Waals radius as defined in NACCESS (45) is used, 100 sampling points are evenly placed on this sphere through the Fibonacci lattice method (48) and a probe radius of 1.4 Å is used. To calculate the relative accessible surface area (RelASA) of a residue, the accessible surface of a residue (ASA) is divided by its maximum accessible surface (MaxASA) as defined in NACCESS (45):

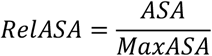

### Conservation determination in Int&in

The conservation is calculated by first using HMMER (3.3) homologous sequences in the Uniref90 database (version 2021_03) (49, 50). Only entries with an E-value≤0.0001, an alignment length of ≥ 70 and a minimum of 35% sequence identity with the input sequence portion are kept to filter out false positives or too small sequences. If multiple domains are found, they are considered as unique hits and subjected to the same filters as described above. If there are more than 2000 sequences, due to technical limitations of the multiple sequence alignment software, these are sampled as follows: first, the number of total sequences is divided by 2000, then the resulting number is iteratively added to itself and the sums, rounded down, are used to sample the sequences. For instance, if 2500 sequences are found, 2500/2000 gives an interval of 1.25, leading to the following sequences being selected: 1, 2, 3, 4, 6, 7, 8, 9, 11,… .This procedure enables reproducible results while sampling through the whole HMMER output. The resulting sequences are then used to create a multiple sequence alignment (MSA) with the sequence in the .pdf file submitted by the user with the muscle program (51). The resulting MSA is then used to create clusters of sequences with ≥ 95% similarity (blank spaces in the MSA are not considered) between each other. One representative sequence out of each cluster is then taken for further evaluation. Due to program-related restrictions, the clusters are once again sampled according to the same principles described above to a maximum of 150 cluster sequences. For inteins, in most cases there are no more than 150 clusters, thus sampling does not occur. The clustered sequences are then passed to rate4site (52, 53), which calculates conservation scores. The conservation scores are then normalized to a scale of 0 and 1 according to the following formulas, similarly to how the bins are created in the ConSurf web-server (54):

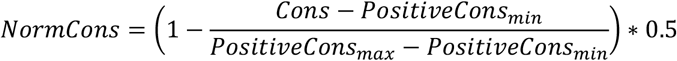

for sites that have a positive conservation (non-conserved sites), with NormCons being the normalized conservation, Cons being the current residues’ conservation, PositiveCons_min_ being the minimal positive conservation out of all residues, and PositiveCons_max_ being the maximal positive conservation of all residues and

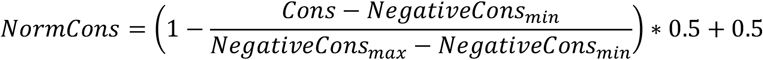

for sites that have a negative conservation (conserved sites), with NormCons being the normalized conservation, Cons being the current residues’ conservation, NegativeCons_min_ being the minimal negative conservation out of all residues, and NegativeCons_max_ being the maximal negative conservation of all residues.

### Binding affinity determination in Int&in

The binding affinity between two fragments is calculated through the PRODIGY prediction model established by Vangone and Bonvin (29, 55). The model is implemented in the backend of the Int&in web server by refactoring the code from Python to C#. The surface accessibility is calculated through an implementation of the algorithm by Lee & Richards (56, 57) as outlined in the FreeSASA source code, with 20 sphere slices and the Van Der Waals radii and maximally accessible surface areas as defined by the NACCESS program (45).

### Secondary structure determination in Int&in

The secondary structure of a sequence is calculated with the DSSP program (58, 59). A split site is considered to be in a secondary structure if both flanking residues are part of a secondary structure.

### Computational resources needed for Int&in

The Int&in web server consists of two distinct parts: a backend written in C# (.Net 3.1) and Python 3 performing the calculations, and a web GUI written in PHP and JavaScript. After submitting a file, the file is stored for at least 30 days on the server and is accessible through a personalized unique key. The file as well as the user’ options are passed from the web GUI to the backend to perform the calculations. The current job status can be accessed through the personalized unique key given to the user. If the user provided an email at submission, a notification will be sent when the job has finished. Otherwise, they will have to check the personalized link to see if the job has finished. The backend makes use of the following libraries: DotNetZip (https://github.com/haf/DotNetZip.Semverd), MailKit (https://github.com/jstedfast/MailKit) and MimeKit (https://github.com/jstedfast/MimeKit). Additionally, the following programs are used for the indicated tasks:

- PDB2PQR v2.1.1 – to generate the structure at a pH of 7 and to add hydrogens
- DSSP (3.0.0) – to generate the secondary structure
- HMMER (3.3) – to identify homologous proteins in the UniRef90 database (2021_03)
- Muscle (v3.8.1551) – to create a multiple sequence alignment (MSA) of the sequence contained in the .pdb file submitted by the user and maximum 2000 sequences extracted from the HMMER output
- Rate4Site (3.0.0.) – to generate the conservation score out of maximum 150 clustered sequences from the MSA
- Python, with the following two libraries: scikit-learn (1.1.0) and pandas

The web-based GUI makes use of the following JavaScript libraries: jQuery (https://github.com/jquery/jquery), NGL viewer for molecular representation (https://doi.org/10.1093%2Fnar%2Fgkv402) and dygraphs for chart plots (https://github.com/danvk/dygraphs). The web-based GUI also lets the user create an account to have an overview of the submitted jobs. The account information is stored in a MYSQL database. Passwords are hashed for security.

### Property significance tests

To assess if the difference between the group of split sites predicted to be active and that of split sites predicted to be inactive was significant when using a defined property, we applied either the t-test or the Mann–Whitney U test (scipy.stats.ttest_ind, mannwhitneyu), depending on whether the groups were considered normally distributed or not, respectively. Normal distribution was evaluated using the Shapiro-Wilk test, the normaltest function from scipy and the *Anderson*-Darling test (scipy.stats.shapiro, normaltest, anderson).

### Property optimization with ANOVA F-Scores

The ANOVA F-Scores were calculated with the sklearn library in Phyton(sklearn.feature_selection.SelectKBest, f_classif).

### Feature combination selection

Different combinations of features were evaluated on the gaussian Naïve Bayes model (from sklearn.naive_bayes.GaussianNB) by calculating the 10x Cross-validated (sklearn.model_selection.KFold with shuffle set to true) Matthews correlation coefficient (MCC, sklearn.metrics.matthews_corrcoef), which was then averaged over ten runs with different seeds (from numpy.random.seed, seeds 0-9 were used).

### Model creation

The gaussian Naïve Bayes model (from sklearn.naive_bayes.GaussianNB; https://jmlr.org/papers/v12/pedregosa11a.html) was trained in Python on the training data set. To evaluate the model in terms of accuracy, precision, recall, and MCC confusion matrixes were generated (sklearn.metrics.accuracy_score, precision_score, recall_score, matthews_corrcoef, confusion_matrix). The confusion matrix representation was generated through seaborn (60). The model was saved through the pickle library in Python.

### Protein structures

The crystal structures of gp41-1 and *Npu* DnaE (PDB id: 6QAZ and PDB id: 4KL5, respectively) were used. The structures were additionally modified in PyMol to remove any extein sequences and mutate residues so that they would represent the native structures. For the gp41-1 intein structure, the first three extein residues (SGG) were removed and the fourth alanine (which inactivates the intein) was mutated back to cysteine. For *Npu* DnaE chain A was used for the computational experiments and the first three extein residues were also removed and the fourth alanine residue was mutated to cysteine; additionally, the last four extein residues (ADNG) in the structure file were also removed. CL as well as all other inteins from the literature data set were modelled through the ColabFold implementation (22) of Alphafold 2 (21).

### Plasmid construction

For the expression of all intein constructs in *E. coli*, the pTrc99A vector was used. A list of all used primers and amino acid sequences of exemplary plasmids is given in *Appendix*, File S5. For PCR amplification the Phusion Flash High-Fidelity PCR Master Mix (2x) from ThermoScientific and the Biometra TOne 96G thermocycler from analytikjena were used. Plasmids were constructed using the NEBuilder® HiFi DNA Assembly Cloning Kit from New England Biolabs.

The *mbp* gene coding for the maltose binding protein (MBP) was amplified from pETM41 via PCR with primers pTrc_MBP_fw and MBP_gp41_rev. Note that the gene in the final plasmids contains a mutation in the first base of the second codon (from A→G), which means that the protein has lysine instead of glutamic acid at that position. Since the anti-MBP antibody worked well we decided against removing the mutation. The *trx* gene coding for thioredoxin was amplified from pETTrx with primers TRX_fw and TRX_SUMO_rev. Plasmids pETM41 and pETTrx were a kind gift of Gunter Stier (Heidelberg University). The *Saccharomyces cerevisiae smt3* gene coding for SUMO was amplified from pTB324_AmiB (kind gift of Thomas Bernhardt, Harvard Medical School, Boston) with primers SUMO_fw and SUMO_rev. The N-fragment of the gp41-1 gene was amplified with primers Gp41-1_fw and Gp41-1_N_merge_rev and the C-fragment of the gp41-1 gene with primer Gp41-1_rev and Gp41-1_C_merge_fw both from pSiMPlk (5). The backbone fragments were amplified with primers pTrc_BB_rev and CoLE1_fw as well as CoLE1_rev and SUMO_BB_fw from pTrc99A in order to have shorter fragments. All fragments (backbone 1, backbone 2, mbp, gp41-1 N-fragment, gp41-1 C-fragment, trx, sumo) contain overhangs allowing for assembly yielding pTRc-MBP-gp41-1-TRX-GS-SUMO-FLAG. The GS linkers and FLAG tags were contained in the respective primers.

Subsequently, GS linkers (GGGGSGGGGS) were added upstream of the local exteins of the N-intein and downstream of the local exteins of the C-intein in pTRc-MBP-gp41-1-TRX-GS-SUMO-FLAG plasmid with primers MBP_GS_rev and CoLE1_fw, Intein_GS_SUMO_fw and CoLE1_rev as well as GP41_GS_Nextein_fw and GP41_Cextein_rev yielding pTRc-MBP-GS-gp41-1-GS-TRX-GS-SUMO-FLAG. The constructs with *Npu* DnaE and CL were constructed using pTRc-MBP-gp41-1-TRX-GS-SUMO-FLAG.

The gene coding for full-length *Npu* DnaE was amplified with primers MBP_GS_rev and CoLE1_fw, Intein_GS_SUMO_fw and CoLE1_rev as well as Npu_CExtein_rev and Npu_NExtein_fw (which also contained a GS linker and the native extein sequences and the exteins) yielding pTRc-MBP-GS-NpuDnaE-GS-TRX-GS-SUMO-FLAG. The gene coding for CL was amplified from pTT32 and pTT43 (25). Specifically, the N-intein was amplified from pTT32 with primers AesN_GS_fw and AesN_rev, while the C-intein was amplified from pTT43 with primers AesC_fw and AesC_GS_rev. The two backbone fragments were amplified from pTRc-MBP-GS-gp41-1-GS-TRX-GS-SUMO-FLAG with primers MBP_GS_rev and CoLE1_fw, Intein_GS_SUMO_fw and CoLE1_rev. The four fragments were then assembled into pTRc-MBP-GS-CSIntein-GS-TRX-GS-SUMO-FLAG.

pTRc-MBP-GS-GS-TRX-GS-SUMO-FLAG containing the exteins and the native local extein sequences for gp41-1 (positive control) was generated from pTRc-MBP-GS-gp41-1-GS-TRX-GS-SUMO-FLAG using primers GS_GP_N_and_C_Ext_fw, CoLE1_rev and GS_Next_rev and CoLE1_fw.

To clone the different split sites, we generated a so-called ‘split cassette’, containing a stop codon, a frame-shifted stop codon followed by a random spacer DNA, a ribosome binding site (61) and a start codon. This split cassette was inserted in the intein-containing plasmid via two overlapping primers encoding the split cassette in their overhangs and annealing to the sequence of the intein at the respective split site (See *Appendix*, File S5 for a list of representative and unique plasmid sequences and a full list of primers. The split cassette primers contain the name of the respective intein and the split site). Two primers annealing to the backbone (CoLE1_fw and CoLE1_rev) were used to generate two fragments with the primers containing the split cassette.

### Bacterial cell lysis

Individual colonies were picked and used to start overnight (ON) cultures, which were grown at 37 °C with 250 rpm shaking in the multitron pro incubator (Infors AG). The next morning, a volume of (100/OD)*3 μl of the ON culture was used to inoculate a fresh tube with 3 ml LB medium plus ampicillin (100 mg/l). The tubes were subsequently shaken at 37 °C and 250 rpm for 90 minutes, after which 3 μl of 1M Isopropyl β-d-1-thiogalactopyranoside (IPTG) were added to each tube. The tubes were then put again inside the incubator and shook for 2 hours and 30 minutes at 37 °C 250 rpm. Afterwards, the OD of each culture was measured and a volume of 100/OD μl was taken and centrifuged at 13.000 rpm for 4 minutes at room temperature. The supernatant was removed from the samples, and the pellets resuspended in 20 μl 4x Lämli Buffer (Bio-Rad) and 80 μl ddH_2_O. The tubes were heated up for 10 minutes at 95 °C and stored at -20 °C. The optical density (OD) of the bacterial cultures was measured with the OD600 DiluPhotometer from IMPLEN GmbH.

### Western Blot

1.5 μl of each cell extract were loaded in a well of a Mini-PROTEAN® 10% TGX(tm) Precast Gel (10 wells, 50 μl pocket volume; Bio-Rad) and separated in the Mini-PROTEAN Tetra Vertical Electrophoresis Cell (Bio-Rad) at 100V for around one hour and 30 minutes. Protein transfer onto a PVDF membrane was carried out with the Trans-Blot Turbo Mini 0.2 μm PVDF Transfer Packs (Bio-Rad) and Trans-Blot Turbo Transfer System (Bio-Rad). The membrane was blocked for 2 hours with 5% BSA-PBST solution at room temperature on a rocking machine at 25 rpm. The blocking buffer was removed and 10 ml 5% BSA-PBST with 1 μl RABBIT anti-DYKDDDDK Tag antibody (Bio-Rad, # AHP1074), 1 μl anti-MBP Monoclonal Antibody (New England Biolabs, #E8032L) and 6 μl anti-*E. coli* RNA Polymerase β antibody (BioLegend, #663905) were added. The membrane was then placed on a rocking machine at 25 rpm for another 2 hours. The BSA-PBST antibody mixture was then removed and the membrane was washed three times with PBST with gentle rocking for 5 minutes. 4 μl of the Cy5 goat anti-rabbit antibody (Invitrogen, #A10523) and 4 μl of the Alexa Fluor goat anti-mouse antibody (Invitrogen, #A11029) were added to 10 ml BSA-PBST and poured over the membrane in a closed box (used to protect it from light). The box containing the membrane was placed on the rocking machine at 25 rpm for another hour. Subsequently, the secondary antibody mixture was removed followed by three washes with PBST, during which the membrane was rocked for 5 minutes in the dark box. The membranes were imaged with the Amersham Typhoon 5 (Global Life Sciences Solutions) with the Cy2 and Cy5 emission filters and excitation wavelengths.

### Calculation of splicing efficiencies

ImageJ2 (1.53s) (62) was used to calculate the efficiency of splicing for each split site according to the following formula:

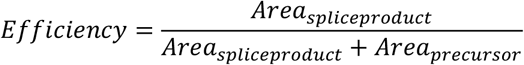

The efficiencies were calculated for both channels (one channel detects the C-extein and one the N-extein). The maximum value was used to train the model.

### Definition of experimentally validated active split site

Two independent WBs were performed for each split site for each tested intein. A split site was considered active only if the splice product was quantifiable in both replicates.

## Acknowledgments

This work was supported by the European Research Council (ERC) under the European Union’s Horizon 2020 research and innovation program (Grant agreement No. 101002044 to BDV), by the Excellence Initiatives of the German Federal and State Governments BIOSS (Centre for Biological Signalling Studies; EXC-294) and CIBSS (Centre for Integrative Biological Signalling Studies; EXC-2189) and partly by the Deutsche Forschungsgemeinschaft (DFG, German Research Foundation) – Project #422681845 within SFB1425.

## Author contributions

MAÖ devised the study. JBB did initial analyses. MS performed experiments, created the algorithm and the website. FT helped with generating split plasmids and the Western blots. JL helped with the Western blots. LX and KV set up the web server. MAÖ supervised the computational work. BDV supervised the experimental work. BDV provided funding and wrote the manuscript with inputs from MS and MAÖ. All authors read and approved of the manuscript.

